# Role of transmembrane helix 6 in substrate recognition of the amino acid transporter MhsT

**DOI:** 10.1101/2020.04.03.022970

**Authors:** Dorota Focht, Caroline Neumann, Joseph Lyons, Ander Eguskiza Bilbao, Rikard Blunck, Lina Malinauskaite, Ilona O. Schwarz, Jonathan A Javitch, Matthias Quick, Poul Nissen

## Abstract

MhsT of *Bacillus halodurans* is a transporter of hydrophobic amino acids and a homologue of the eukaryotic SLC6 family of Na^+^-dependent symporters for amino acids, neurotransmitters, osmolytes, or creatine. The broad range of transported amino acids by MhsT prompted the investigation of the substrate recognition mechanism. Here, we report six new substrate-bound structures of MhsT, which, in conjunction with functional studies, reveal how the flexibility of a Gly-Met-Gly (GMG) motif in the unwound region of transmembrane segment 6 (TM6) is central for the recognition of substrates of different size by tailoring the binding site shape and volume. MhsT mutants, harboring substitutions within the unwound GMG loop and substrate binding pocket that mimick the binding sites of eukaryotic SLC6A18/B^0^AT3 and SLC6A19/B^0^AT1 transporters of neutral amino acids, exhibited impaired transport of aromatic amino acids that require a large binding site volume. Conservation of a general (G/A/C)ΦG motif among eukaryotic members of SLC6 family suggests a role for this loop in a common mechanism for substrate recognition and translocation by SLC6 transporters of broad substrate specificity.

## Introduction

The solute carrier 6 (SLC6) subfamily is part of a larger amino acid-polyamine-organocation (APC) transporter superfamily [1] [2]. Twenty different human genes encode SLC6 transporters that are responsible for the active transport of a variety of solutes including neurotransmitters such as serotonin, dopamine and norepinephrine, as well as creatine, taurine, choline, betaine and amino acids [1]. Amino acid transporters of the SLC6 family, such as the γ-aminobutyric acid transporter (GABA transporter, GAT), glycine transporters (GlyT) and the neutral amino acid transporters SLC6A18 and SLC6A19 participate in the active reuptake of amino acids in kidneys, small intestine [3] and brain tissue [4, 5]. Malfunctions of SLC6 transporters are associated with a number of neurological and metabolic diseases, such as schizophrenia [6], epilepsy [7, 8], depression [9, 10] [11] and aminoacidurias [12, 13]. Loss-of-function mutations of the neutral amino acid transporter SLC6A19 (B^0^AT1), the major amino acid uptake system in the gut, are causative of Hartnup disorder, where amino acid uptake is insufficient and in particular affecting tryptophan levels and therefore biosynthesis of niacin, melatonin, and serotonin [14].

Structural characterization of eukaryotic SLC6 proteins has been based on crystal structures of the dopamine transporter from *Drosophila melanogaster* (dDAT) [15] and the human serotonin transporter (hSERT) [16]. They both belong to the neurotransmitter:sodium symporter (NSS) family and feature the so-called LeuT fold [17] that was first identified in the crystal structure of the NSS homologue LeuT from *Aquifex aeolicus* [18].

The LeuT fold displays an inverted pseudo-two-fold symmetry between two transmembrane helix (TM) bundles that is conserved in many other families of symporters and exchangers [19–24]. In the LeuT structure the primary substrate (S1) binding site and the two sodium ion (Na1 and Na2) binding sites are located between the so-called bundle and scaffold domains (Figure 1A and B). Proposed first for LeuT [25], a second, allosteric substrate site S2 has been associated with an extracellular vestibule of the transporter [26, 27]. It has not been possible to capture crystal structures of MhsT or LeuT in S2-bound states, which from single-molecule studies appear dynamic in nature [27], but inhibitor binding at an overlapping site has been observed [15, 28–32].

**Figure 1:**
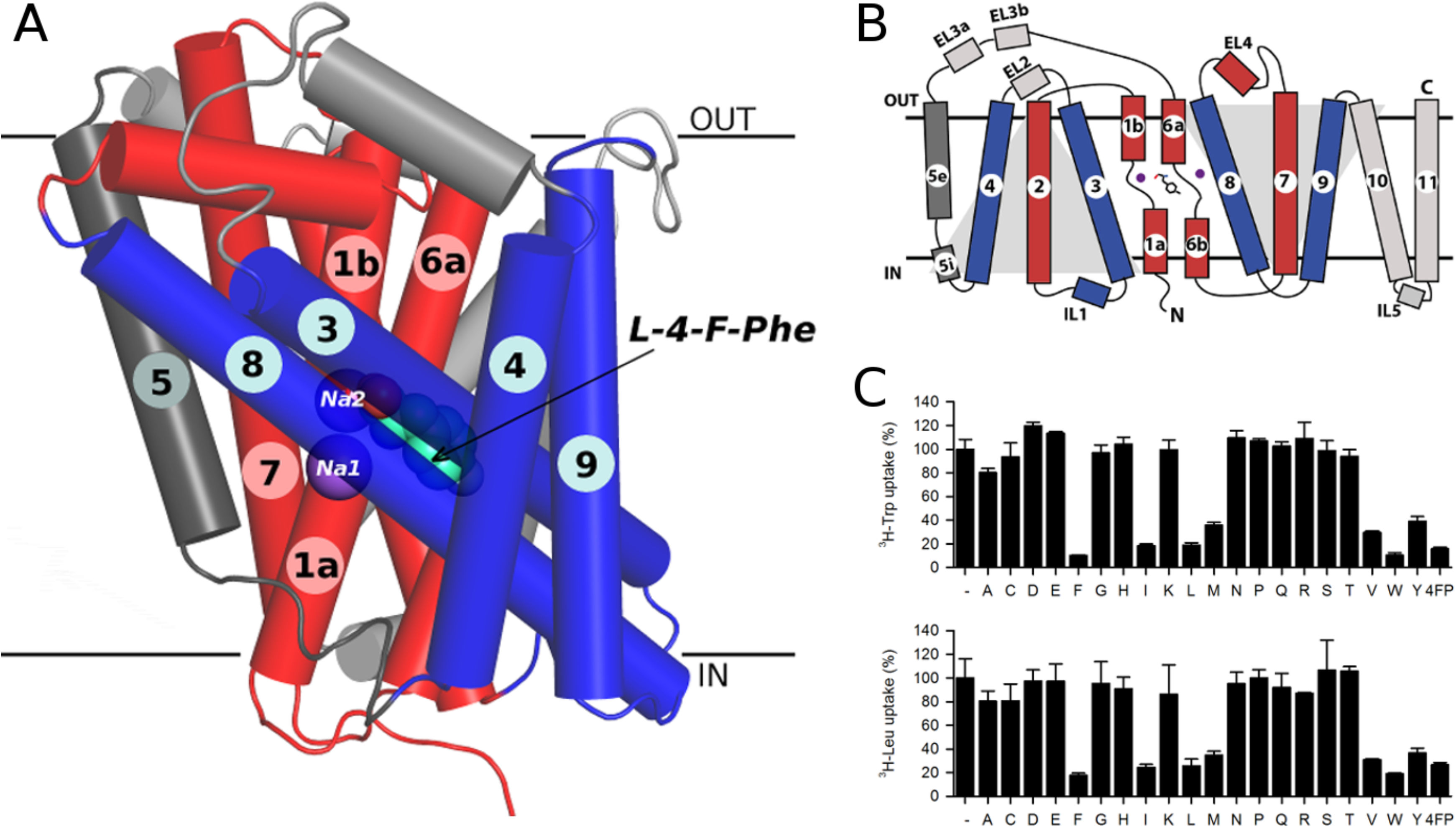
Overview of MhsT structure and function. A) MhsT is shown in a ribbon representation (scaffold helices in grey, bundle helices in red and blue, and TM5 in dark grey, sodium ions shown as purple spheres, L-4-Fluoro-phenylalanine, bound between unwound region of TM1 and TM6, shown as grey spheres. B) Schematic representation of the MhsT structure (LeuT fold). Color coding same as in panel A. C) Results of a competitive uptake assay showing inhibition of 0.2 μM ^3^H-Trp or ^3^H-Leu uptake in MhsT-WT-expressing MQ614 cells, measured for 30 seconds in the presence of 12 μM hydrophobic amino acids.

We have reported structures of another SLC6 orthologue, MhsT (multiple hydrophobic amino-acid substrate transporter) from *Bacillus halodurans,* revealing an occluded, inward-oriented state with bound Na^+^ and L-tryptophan in the Na1 and Na2 sites and the substrate site S1, respectively [33]. It was previously reported that MhsT displays a broad specificity for hydrophobic amino acids [33, 34] (Figure 1C). Interestingly, based on sequence conservation, MhsT is a close orthologue of the human SLC6A18 and SLC6A19 (also known as B^0^AT3 and B^0^AT1 with sequence identities of 36% and 30%, respectively). These are major amino acid transporting systems for non-polar amino acids in the brush border membrane of epithelial cells [4] in kidney proximal tubules [3, 35] and intestine (jejunum) [3]. SLC6A19 was reported to facilitate Na^+^-dependent transport of its substrates (L-Leu, L-Ile, L-Val, L-Met, L-Phe, L-Trp, L-Thr, and L-His) with millimolar apparent affinities [14]. SLC6A18 exhibits 50% sequence identity to SLC6A19 and, to some extent, an overlapping substrate specificity. It has been shown to transport aliphatic amino acids (L-Ala, L-Met, L-Val, L-Ile, L-Gly, L-Ser, L-Leu) also with millimolar apparent affinities [36].

Here we report crystal structures of MhsT complexed with six different substrates, exploiting further the inward-oriented, occluded state, in an effort to identify the structural elements of the S1 binding site underlying substrate recognition, promiscuity, and transport. We find that the architecture of the S1 site is broadly defined by two possible states that are dependent on the nature of the hydrophobic amino acid substrate: smaller aliphatic amino acids (L-Val, L-Leu and L-Ile) and larger aromatic amino acids (L-Trp, L-Phe, L-Tyr and the L-Tyr analogue L-4-F-Phe). This modulation centers on a key role of the unwound segment of TM6 that defines the binding site shape and volume and in both cases compatible with a flexible transport mechanism. Additionally, we used site-directed mutagenesis to investigate the role of various residues involved in substrate binding and their influence on substrate specificity and transport.

## Results

### MhsT crystallization, processing and refinement

Previously, crystals of MhsT bound to L-Trp and Na^+^ were obtained using high concentrations of lipid and detergent (HiLiDe) [37] and the lipid cubic phase (LCP) [38] methods. Using the HiLiDe crystallization conditions, we have obtained six structures of MhsT in complex with six different ligands in the S1 site (Figure 2 and Figure S6): the tyrosine analog 4-fluoro-L-phenylalanine (4F-Phe), L-Tyr, L-Phe, L-Ile, L-Leu and L-Val. Protein-substrate crystals (Figure S1) were obtained in similar conditions to those previously reported [33].

**Figure 2:**
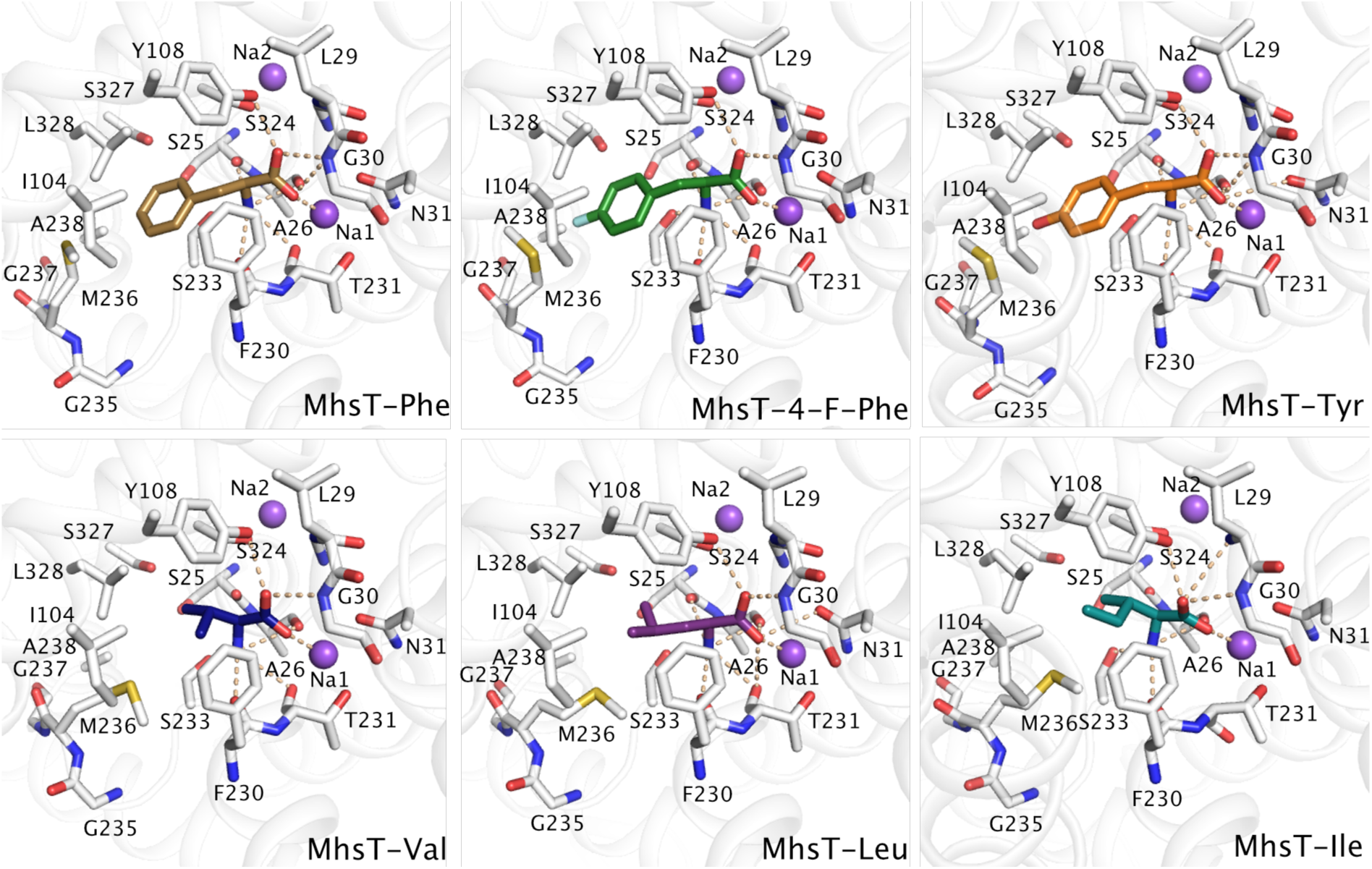
Comparison of the ligand binding sites of MhsT in complex with the various substrates: L-Phe, L-4F-Phe (Tyr anlogue), L-Tyr, L-Val, L-Leu and L-Ile. Protein is shown as white ribbon with the binding site defined with white sticks. Substrates are visualised as coloured sticks and the sodium ions as purple spheres. Hydrogen and electrostatic interactions are identified by broken lines.

Structures of MhsT in complex with 4F-Phe, Tyr and Phe were determined at about 2.3 Å resolution (Table 1) in the P2 space group with one molecule in the asymmetric unit similar to the Trp-bound complex [33]. The complexes with Leu and Val substrates crystallized in a different P2_1_ crystal form with two molecules in the asymmetric unit related by twofold NCS, and their structures were determined at 2.35 Å and 2.60 Å resolution, respectively (Table 1). The dataset for MhsT-Leu showed pseudomerohedral twinning with a twin fraction close to 0.5 and therefore was refined using the appropriate twin law (h, -k, -l) resulting in a large drop of R-factors. The dataset for MhsT-Val exhibited no notable twinning. The MhsT-Ile complex crystallized in a distinct P2_1_ crystal form with translational non-crystallographic symmetry (tNCS) that exhibited strong radiation sensitivity and crystal-to-crystal nonisomorphism, but a dataset with completeness of 80% at 3.1 Å resolution could be obtained by merging of data collected from two crystals (Table 1). The presence of translational NCS in this crystal form combined with the limited completeness of the crystallographic data impaired refinement, but the structure determination was sufficiently clear to discern important features (see below).

**Table 1:**
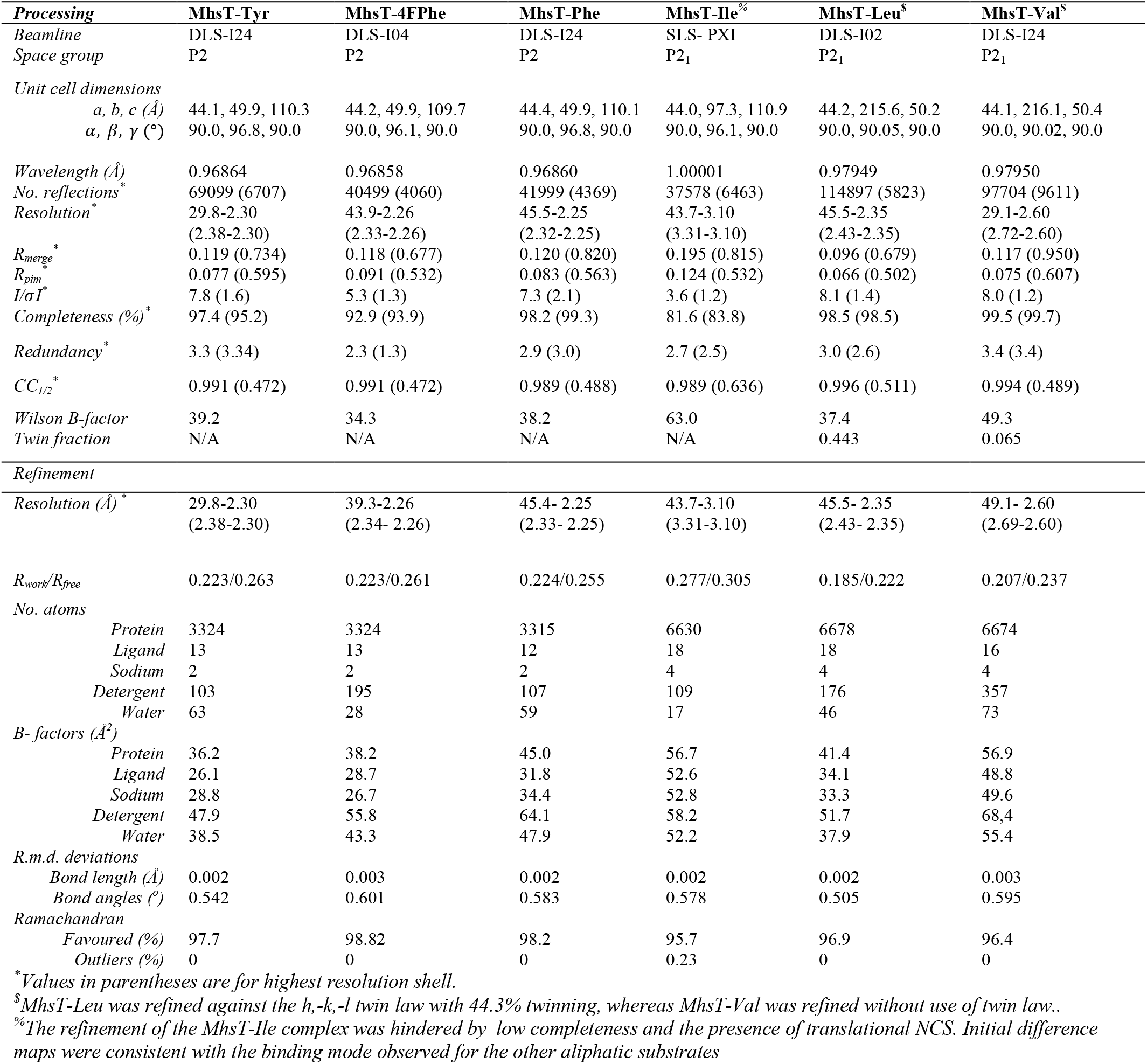
Data reduction and refinement statistics.

### Substrate-bound MhsT

All crystal structures of MhsT captured the protein in an inward-facing occluded state, with a closed extracellular vestibule, an ordered N-terminal tail associated to the intracellular surface, and an unwound TM5 (Figure 1A) within a conserved ProX_9_Gly motif as previously described [33]. This state allows initial solvation of the Na^+^ ion at the Na2 site from the cytoplasmic environment. The various substrate-bound structures are overall similar and superimpose with a low *Cα* r.m.s.d. (Table 2). However, local differences are observed that allow the S1 site to accommodate substrates of different size/volume.

**Table 2:** r.m.s.d. values of the different complexes when compared to MhsT-Trp (PDB ID: 4US3) and comparison of binding site cavity volumes for the substrate bound MhsT structures.

| Complex | r.m.s.d. value | Cavity volume on plot grid ($\text{\AA}^3$)* |
| --- | --- | --- |
| **MhsT-Trp** | n.a. | 230 |
| **MhsT-Tyr** | 0.185 | 224 |
| **MhsT-4FPhe** | 0.165 | 215 |
| **MhsT-Phe** | 0.162 | 210 |
| **MhsT-Ile** | Chain A : 0.351 | 159 |
|  | Chain B : 0.340 |  |
| **MhsT-Leu** | Chain A : 0.292 | 158 |
|  | Chain B : 0.287 |  |
| **MhsT-Val** | Chain A: 0.274 | 144 |
|  | Chain B: 0.267 |  |

\*Calculated with voidoo software [65]

The substrates are bound with the amino acid moiety within the S1 site similar to the MhsT-Trp structure [33] (Figure 2, Figure S3 and Figure S6). The substrate amino group forms hydrogen bonds with backbone amide oxygen atoms of Ala26 in TM1, Phe230, Thr231 and the side chain of Ser233 in TM6, while the carboxyl group interacts via hydrogen bonds to the side chain of Tyr108, the backbone amide nitrogen of Gly30 and Na^+^ in the Na1 site. Additionally, the positive dipole on TM1b interacts with the negatively charged carboxyl group of the substrate, whereas the negative dipoles of TM1a and TM6a point towards the positively charged amino group.

The side chains of hydrophobic substrates are located to a separate pocket defined by the side chains of Ile104 in TM3, Phe229, Phe230, Ser233, Met236, Ala238 in TM6, Val331, Ser324, Ser327 and Leu328 in TM8, Leu393 in TM10 and the backbone atoms of Ala26-Leu29, Phe229, Thr231 and Leu324. The hydrophobic nature of the side chain binding pocket explains the clear preference of MhsT for hydrophobic substrates over polar/charged amino acids (Fig. 2).

### Discrimination of aromatic and aliphatic amino acid binding modes

The different substrate complexes of MhsT illustrate the changes required for the substrate binding pocket to accommodate the disparately sized substrates. Although the seven substrates (including L-Trp-bound structure; PDB ID: 4US3) bind the S1 pocket in a similar mode concerning the amino acid group, significant structural changes within the hydrophobic cavity are observed that serve to modulate the S1 site on the basis of the nature and size of the substrate’s side chain, i.e., aliphatic or aromatic.

Binding of aromatic amino acids was characterized using MhsT structures with bound 4F-Phe, Tyr, Phe (this study) and Trp [33]. A comparison of these four structures show a small conformational change of the rotamer of Met236 in MhsT-4F-Phe and MhsT-Tyr compared to the two other structures (Figure 3A). For the 4F-Phe and Tyr complexes, the sulphur atom of Met236 is pointing in the direction of the fluorine atom of 4-F-Phe making a polar interaction, and the hydroxyl group of Tyr making a favorable hydrogen bond, which is not possible for Phe and Trp substrates, where instead the hydrophobic methyl group of Met236 points in the direction of the ligands for hydrophobic interactions.

**Figure 3:**
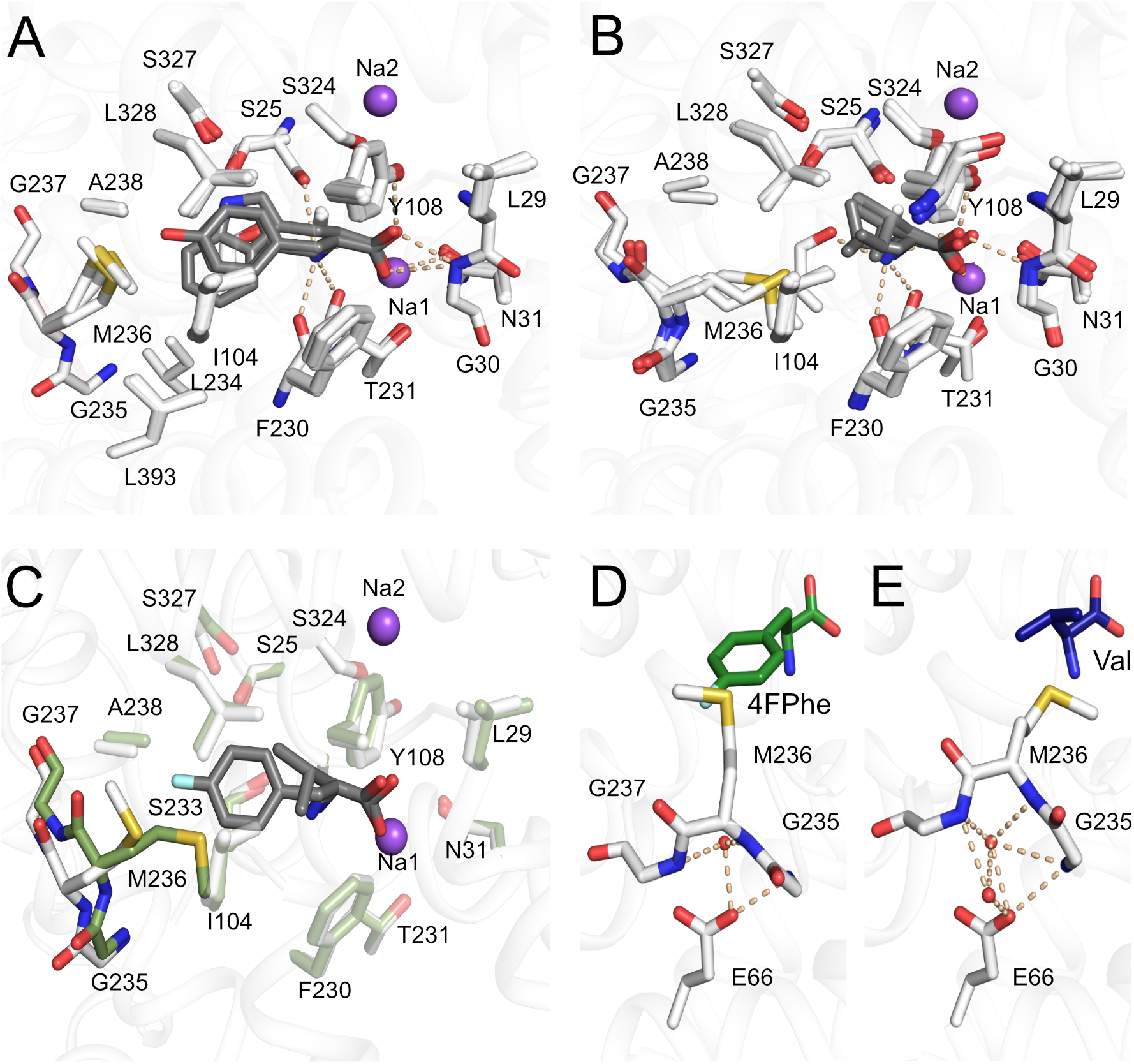
Grouping of binding sites: A) aromatic ligands consisting of L-Trp, L-Phe, L-4-F-Phe, L-Tyr, B) aliphatic ligands consisting of L-Val, L-Leu and L-Ile. C) Overlay of MhsT-Val and MhsT-4FPhe structures to visualize changes upon binding of different sized ligands. The unwound region of TM6 is non-transparent. D) Conformation of the unwound part of TM6 in case of an aromatic substrate (L-4-F-Phe as an example) and E) aliphatic substrate (L-Val as an example). Glu66 is coordinating the GMG loop through water molecules, in the case of aromatic substrates only one water molecule is found, whereas in the case of aliphatic substrates two water molecules are present. Protein is shown as white ribbon with the binding site defined with white sticks. Substrates are visualised as grey sticks and the sodium ions as purple spheres.

Binding of the aliphatic amino acids was characterized using structures obtained for the MhsT-Val, MhsT-Leu and MhsT-Ile complexes (Figure 3B). Comparison of aliphatic and aromatic substrate complexes highlighted that the non-helical fragment of TM6, formed by residues Leu234-Gly235-Met236-Gly237-Ala238, changes its relative position within the binding site depending on the size of the ligand (Figure 3C and Figure S7). This loop will be referred to as the GMG motif, as the shift of these three residues is most significant. The movement is most pronounced when the MhsT-Val complex is compared to the MhsT-Trp structure. The presence of a small substrate in the binding site prompts the inward movement of the GMG loop thus compensating for the smaller substrate side chains. The unwound region of TM6 is displaced by approx. 2 Å toward the bound substrate when comparing the position of Met236 Cα. The inward movement of the GMG loop reduces the volume of the binding pocket significantly, with volumes approaching 230 Å^3^ for the aromatic substrates and diminishing to 144 Å^3^ in the case of valine (Table 2, Figure 4, Figure S3 and Figure S4). Taken together the seven substrate-bound structures describe a bi-modal binding site that discriminates between apolar aliphatic and aromatic amino acids by a main chain movement of the GMG motif, and by finetuning of the position of individual side chains through rotamer changes of the M236 side chain.

**Figure 4:**
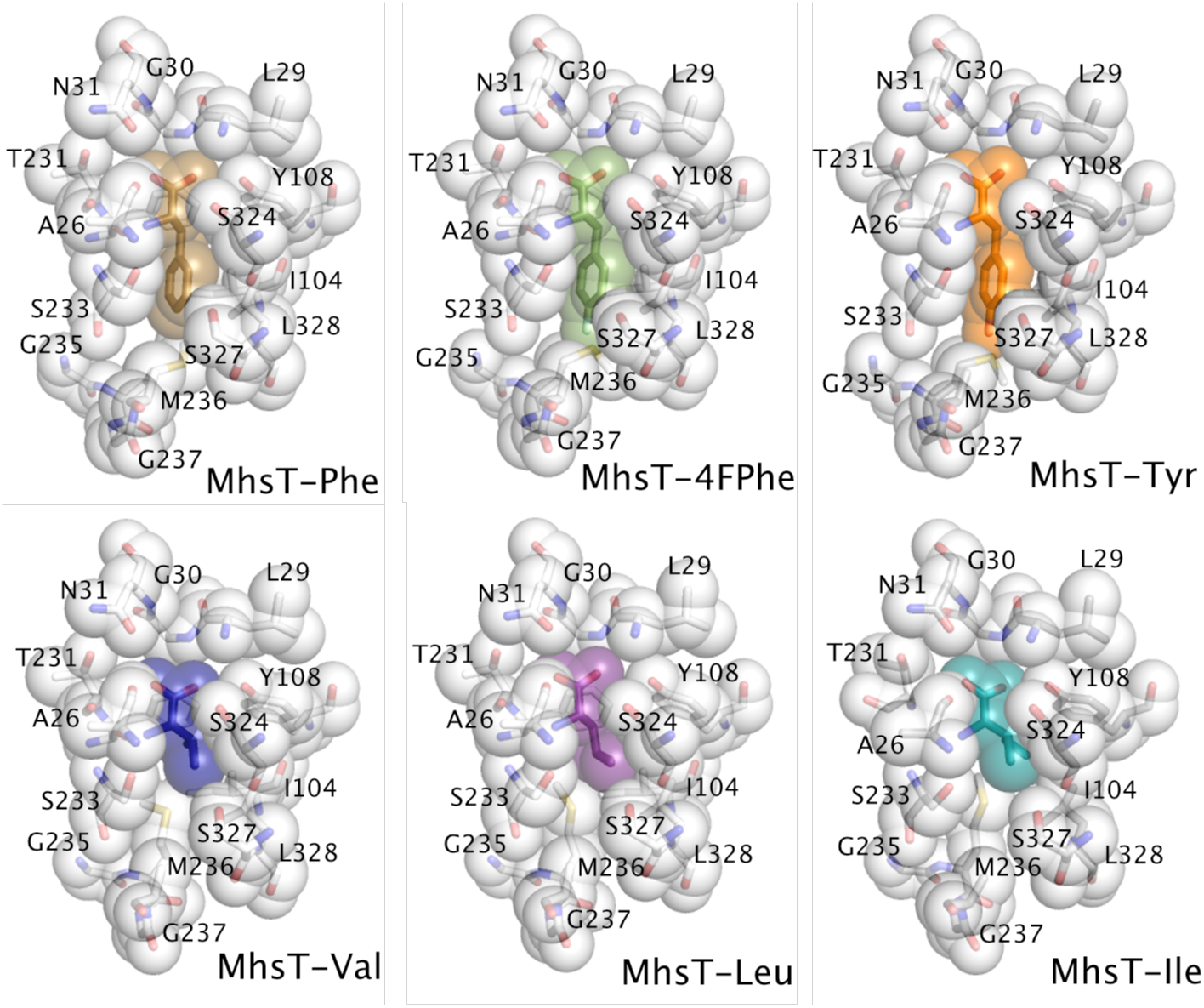
Volumes of the binding pockets visualised by mean of the van der Waals spheres of the ligand and coordinating residues of MhsT in complex with L-Phe, L-4-F-Phe, L-Tyr, L-Val, L-Leu and L-Ile.

Overall changes in the volume of the MhsT hydrophobic cavity upon binding of different substrates follows an “induced-fit” mechanism [39, 40]. Specifically the hydrophobic nature of the substrate and its binding pocket disfavors the possibility of increased solvation compensating for the deficit in substrate volume, thus promoting the intrinsic structural remodeling of the substrate binding site – a mechanism, which is also reminiscent of the movement of the unwound segment of TM1 to compensate the empty hydrophobic-lined substrate binding site in the occluded return state of LeuT [41].

### The conserved Glu66

Proximal to the substrate binding site, however, a glutamate residue (Glu66) is buried in the MhsT structure and interacts with the backbone of the GMG motif of TM6. Interestingly, Glu66^MhsT^ is conserved throughout the SLC6 family (Figure S2A). Mutagenesis of the equivalent glutamate residue in SERT [42], DAT [43, 44], NET [45] and GAT1 [46] diminishes transport activity, supporting its important role in substrate translocation. In SERT, interaction between this glutamate (Glu136^SERT^) and TM6 has been proposed to be crucial for conformational transitions of the protein [42], allowing for changes between the outward and inward facing states associated with the transport cycle. MhsT offers the opportunity to analyse interactions of this residue in the occluded inward-oriented state.

Even though the unwound region of TM6 adopts different conformations in the aliphatic and aromatic substrate bound MhsT complexes, interactions with Glu66 are maintained. These interactions proceed through both direct and more flexible water mediated hydrogen bonds (Figure 3D-E and Figure S8). With smaller aliphatic substrates bound to MhsT, the space formed through the displacement of unwound TM6 (including GMG) is filled with an additional ordered water molecule interacting with both Glu66 and unwound TM6. The water molecules could have two functions i) to stabilize the different conformations of unwound TM6 (B-factor analysis of all complexes revealed that the conformation of unwound TM6 is largely rigid for all bound substrates, Figure 5), and ii) to preserve the interaction with Glu66 and its critical role in transport.

**Figure 5:**
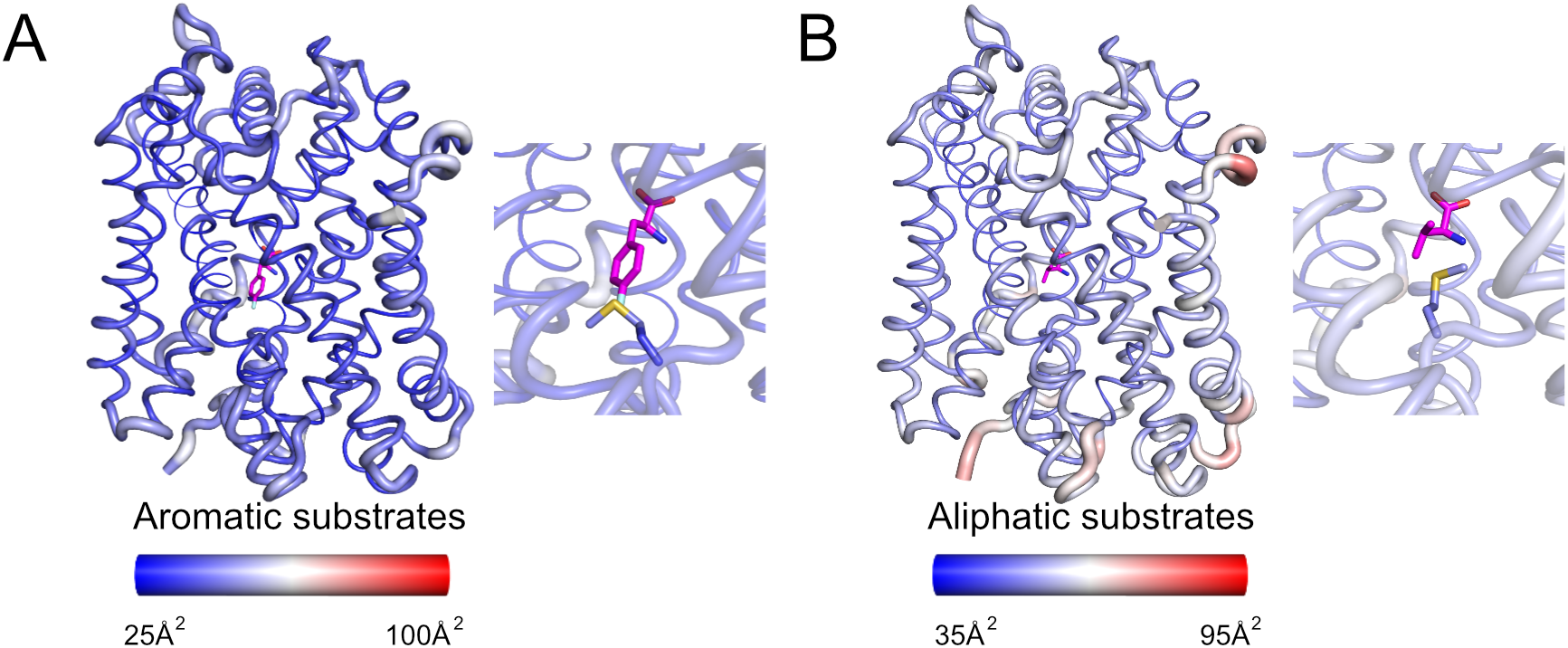
Structural flexibility analysis of MhsT in complex with A) L-4-F-Phe and B) L-Val as representans of the aromatic and aliphatic substrates, respectively, with the ligands represented as magenta sticks. The analysis was visualised by atomic putty and thermal gradient representation of B-factor in Pymol. Red color indicates a high degree of disorder, while blue color indicates a low degree of disorder. In the inset, the substrates as well as the GMG loop are highlighted.

**Figure 6:** A) Specificity of MhsT-M236F. Uptake of 0.1 μM ^3^H-Leu was measured for 10-second periods in the presence or absence of the indicated natural amino acids and 4-fluorophenylalanine (at 100 μM final concentration) in MQ614 expressing MhsT-WT or –M236F. B, C) Time course of 0.1 μM ^3^H-Leu (B) or 0.1 μM ^3^H-Trp (C) by MQ614 expressing MhsT-WT (■) or –M236F (∇), or MQ614 transformed with the control plasmid, pQE60 (◯). D, E) Kinetics of ^3^H-Leu (D) or ^3^H-Trp (E) uptake. Data are the mean ± S.E.M. of triplicate determinations of a representative experiment and the kinetic constants are the mean ± S.E.M of the fit.

Indeed, a comparison of the various known LeuT and MhsT structures in different states highlights a consistent interaction of Glu66 with the unwound TM6 in spite of the significant rearrangements of TM2, TM6a and TM6b during the transport cycle (Figure S5). The interaction is also maintained when different sized substrates are bound, underlying the importance of this interaction. The Glu66 residue could be viewed as a fulcrum at the protein core about which changes related to the transport mechanism happen.

### Conservation of the GMG loop

Unlike the considerable conservation of the unwound region of TM1 in SLC6 transporters, which has recently been reported as important also for the substrate-free return transition [41], the unwound region of TM6 is far more diverse in sequence. The central residue, Met236 in MhsT, is typically a non-polar residue ranging in size from leucine (e.g. GABA transporter) to tryptophan (e.g. glycine transporter) with a majority of SLC6 transporters, like LeuT, having a phenylalanine (e.g. serotonin and dopamine transporters). A C-terminally flanking glycine, Gly237 in MhsT, is fully conserved in the SLC6 family (Figure S2B), while the N-terminally flanking residue, Gly235 in MhsT, is typically a cysteine, glycine or alanine. The flanking residues may determine the degree of flexibility of the loop, which, combined with the diversity of the unwound TM6 sequence, is likely related to the substrate specificity of the transporter, eventually with promiscuity to a broader range of substrates as for the GMG motif of MhsT.

Like MhsT, the closely related amino acid transporters SLC6A18 (B^0^AT3) and SLC6A19 (B^0^AT1) are neutral amino acid transporters, although with a specific preference for aliphatic amino acids [1, 36, 47]. Sequence alignment of MhsT, SLC6A18 and SLC6A19 identified similarities and differences between their binding pockets (defined as residues within 4 Å from bound substrate in MhsT, Table 3). The substrate binding site of MhsT exhibits ~50% conservation compared to SLC6A18 and SLC6A19. In both transporters, the unwound TM6 motif is AFG, with the reduced flexibility and bulkier central Phe residue putatively defining the substrate specificity towards a preference for aliphatic amino acids similar to LeuT. Interestingly, in both SLC6A18 and SLC6A19 the AFG motif precedes an additional glycine that may offset this reduced flexibility and bulkier central residue (Figure S2).

**Table 3:** Comparison of the binding site residues of MhsT, SLC6A18 and SLC6A19.

| **MhsT** | **LeuT** | **SLC6A18** | **SLC6A19** | **dDAT** | **SERT** |
| --- | --- | --- | --- | --- | --- |
| Ala26 | Ala | Ala | Cys | Ala | Ala |
| Leu29 | Leu | Leu | Leu | Leu | Leu |
| Gly30 | Gly | Gly | Gly | Ala | Gly |
| Asn31 | Asn | Asn | Asn | Asn | Asn |
| Ile104 | Val | Ile | Val | Ile | Ile |
| Tyr108 | Tyr | Tyr | Tyr | Tyr | Tyr |
| Phe230 | Phe | Phe | Phe | Phe | Phe |
| Thr231 | The | Ser | Ser | Ser | Ser |
| Ser233 | Ser | Ser | Ser | Gly | Gly |
| Met236 | Phe | Phe | Phe | Phe | Phe |
| Ala238 | Ala | Gly | Gly | Val | Val |
| Ser324 | Ser | Thr | Ser | Ser | Ser |
| Ser327 | Ala | Gly | Gly | Ala | Gly |
| Leu328 | Ile | Thr | Asn | Gly | Gly |
| Val331 | Pro | Ala | Gly | Gly | Ala |

To probe the role of the unwound region of TM6 and the composition of the substrate binding site on substrate specificity of the SLC6 transporters, we generated mutant forms of MhsT in which the binding site mimicks the binding sites of SLC6A18 and SLC6A19. These MhsT mutants were subsequently used for transport studies.

### *In vivo* uptake experiments

A competitive uptake assay of L-[^3^H]Leu and non-radiolabeled amino acids allowed for an investigation of the substrate specificity of the generated MhsT variants. From the group of single mutants, the most striking effect was observed for the M236F variant, which showed complete insensitivity towards the aromatic amino acids (Figure 6A), while the interaction of hydrophobic but non-aromatic substrates seems to be unaffected in this variant. Measurements of initial rates of transport in MQ614 cells expressing MhsT-WT or -M236F confirmed comparable kinetics of the two proteins for Leu (Figure 6). The *V_max_* of Leu transport was calculated to be 3.23 ± 0.14 nmol x min^-1^ x mg of cell protein^-1^ for MhsT-WT and 3.33 ± 0.13 nmol x min^-1^ x mg of cell protein^-1^ for MhsT-M236F. The *K_m_* and *V_max_* of Trp uptake by MhsT-WT were 1.07 ± 0.19 μM and 2.59 ± 0.12 nmol x min^-1^ x mg of cell protein^-1^, respectively, but could not be determined for MhsT-M236F, suggesting complete abolishment of L-Trp uptake by M236F variant (Figure 6C).

## Discussion

The total of seven substrate-bound crystal structures of MhsT, six of them determined here, highlight the structural reorganization, centered on the unwound region of TM6, that is required to accommodate the transport of aliphatic and aromatic residues. This movement tailors the S1 binding site volume and provides a structural basis for the promiscuity of the transporter. A single point mutation of Met236 to Phe, found in ~50% of human SLC6 and also in e.g. LeuT, abolishes the transport of aromatic amino acids by MhsT, but maintains transport of aliphatic side chains.

Comparison of the various substrate-bound MhsT structures illustrates that the GMG motif present in the unwound region of TM6, and conserved to some extent among other bacterial homologues and eukaryotic members of NSS family (Figure S2B), serves as a regulator of the binding site pocket volume and modulates the chemical environment. Involvement of the non-helical fragment of TM6 in substrate binding was also observed in crystal structures of LeuT with different amino acids (L-Leu [17], Gly, L-Ala, L-Met) and their derivatives (L-4-F-Phe [48]). Formation of the LeuT complex with L-Ala and Gly is possible due to torsions of two spatially neighboring residues, Phe259 and Ile359, by about 30° and 15°, respectively, when compared to the L-Leu complex. These two residues were identified as the volumetric sensors in LeuT and mediators of the allosteric communication between the substrates in the S1 site and the intracellular gate [49, 50]. Indeed, the Phe259 residue in LeuT corresponds to Met236 in MhsT, with both of them flanked by Gly residues (Figure S2B). While binding of the various aliphatic substrates by LeuT is associated with the same binding site architecture as for the LeuT-Leu complex, the much bulkier 4F-Phe requires the largest rearrangements of LeuT; Ile359 undergoes a ~180° rotation of its side chain, while the backbone of the unwound region of TM6 is slightly shifted outward in order to accommodate the substrate within the protein cavity [51]. This is similar, but significantly less pronounced, than the outward movement of unwound TM6 region in the MhsT aromatic substrate complexes. Singh *et al* speculated that the displacement of the unwound TM6 region in the LeuT-L-4F-Phe structure marked a strained occluded state, with a reduced rate of transition to the inward-open state, consistent with the lower tyrosine turnover rate [51]. Furthermore, the larger L-Trp acts as a competitive inhibitor for LeuT and traps the transporter in an outward-open conformation. In light of our findings for MhsT, this movement of unwound TM6 in LeuT could also be viewed as an extension of the binding pocket in order to accommodate larger substrate side chains. It is important to note that while both MhsT and LeuT transport a broad range of hydrophobic amino acids, their specificities seem to be shifted towards opposing ends of the spectrum. LeuT, similarly to SLC6A18, is more suitable to the transport of smaller, aliphatic substrates (Gly, L-Ala, L-Leu) with a maximal catalytic efficiency for L-Ala, whereas MhsT, having high apparent affinity for bulky, aromatic ligands (L-Tyr, L-Phe and L-Trp), resembles the activity of the SLC6A19 transporter. It seems that the two transporters use different mechanisms to accommodate for different substrates sizes. In LeuT, changes in the conformation of the binding site residues and a slight movement of the nonhelical part of TM6 are required, whereas MhsT bases its promiscuity in the S1 site largely on a bimodal switch of the structure of the GMG motif.

The movement of the unwound region of TM6 was also observed in crystal structures of the *Drosophila melanogaster* dopamine transporter (dDAT) in complex with substrates like dopamine or amphetamines [52] and inhibitors like antidepressants [15, 29] or tropane inhibitors. In the case of dopamine, metamphetamine, or D-amphetamine, the non-helical fragment of TM6 is moved into the binding pocket in a similar way as for MhsT in complex with aliphatic substrates. The binding of substrates is accompanied by specific conformations of Phe319^dDAT^ (Phe230^MhsT^) and Phe325^dDAT^ (Met236^MhsT^) that additionally reduce the size of the binding pocket. Inhibitors lock dDAT in an outward-open conformation, either by blocking the binding site with multiple aromatic rings, like in the case of antidepressants, where the unwound part of TM6 is moved out and the two abovementioned Phe turn away from the pocket, or by hindering the movement of the extracellular gate as in the case of tropane inhibitors. Here, the unwound part of TM6 and the two Phe residues are in similar conformations as the in the case of the substrates [15].

Since the GMG motif in MhsT and its corresponding loop in LeuT and dDAT can be put into focus as a possible important mechanism for substrate size compensation, it is tempting to speculate whether the flexible structure of the protein regions around the binding site can have conserved function among other transporters. Probably, the structural features, similar to the observed (G/A/C)ΦG motif (GMG for MhsT, GFG for LeuT, and AFG, AWG, CLG, CQG, GFG and GLG for human SLC6 members), would be relevant in transporters that facilitate translocation of a broad range of solutes, especially those of different sizes. Functional studies in *E.coli* cells expressing WT or mutant variants of MhsT, the latter of which mimicking the neutral amino acid transporters SLC6A18 and SLC6A19, revealed loss-of-function to transport aromatic amino acids (Figure S9) with M236F (of the GMG motif) being the most significant in its effect on function of MhsT (Figure 6). Other single mutations had moderate to no effect. Comparable levels of expression of all of the constructs were confirmed suggesting that any alterations of the transport rates were caused by lowered activity of the protein. The observed phenotype can be easily explained by an alteration of the binding site as well as a change within the electrostatic and steric environment of the hydrophobic pocket.

Based on the available crystal structures and competitive uptake assays, we can assume the Val side chain represents the smallest ligand suitable for transport by MhsT-WT (Figure 2 and Figure 6A). The size of Gly or Ala may be too small to be compensated by an inward tilt of the GMG loop, as Met236 presumably cannot stabilize the requirement of a smaller binding site. The M236F substitution did not result in a gain of glycine or alanine transport activity showing that also other features in the S1 site play a role in defining the substrate specificity. Possibly, mutagenesis of several residues, e.g. placed at the interface of more distant TM helices, would be required to alter the overall shape and dynamics of the binding site and thereby its binding specificity along with transport activity. Furthermore, it would be interesting to understand the specifics of the interaction of the different amino acid substrates in the allosteric S2 site in context of the overall mechanism and partial reactions of the transport cycle, but remains obscure in the absence of relevant structures.

In a computational study on LeuT, the binding site residues placed in TM6 (Phe259 of the GFG motif) and TM8 (Ile359) were found to have the highest contributions in communication between the substrate binding site and the intracellular gate of LeuT, thus serving as important residues in allosteric signaling during the substrate transport [49, 50]. Also, among the residues present within the LeuT S1 site, Phe259 was reported to have the highest contribution to coordination of the intracellular gate. In general, the findings highlight TM6 as a major mediator in coordination of the intracellular gate by residues from S1 and S2 sites of LeuT with F259 (MhsT M236) being proposed to serve as a regulatory residue which, e.g. in the case of L-Trp binding to LeuT, leads to inhibition [49].

As observed also in the MhsT-substrate complexes, the unwound region of TM6 interacts with a highly conserved Glu66 from TM2. The residue, when mutated in SERT, was shown to reduce or abolish the transport, while not affecting the substrate binding, supporting the hypothesis that the residue is crucial for conformational transition of the protein [42]. The negative influence of the conserved Glu substitutions was also reported for DAT [43], [44] and NET [45], suggesting its important role in substrate binding and translocation. In fact, the obtained MhsT structures revealed, depending on the size of the bound substrate, the presence of additional water molecules involved in the coordination of the unwound region of TM6 to Glu66 and slightly changed H-binding networks. In the case of all MhsT-substrate complexes, the communication between the GMG loop and the conserved Glu66 from TM2 is maintained both directly and through interactions with additional water molecules (Figure S8). The presence of the water molecules around TM6, and a substrate inside the binding pocket, will subsequently allow for the transporter transition from the substrate-bound, outward facing state towards the inward-facing conformation and substrate release. These water molecules are also found in the LeuT structures [53]. A comparison of the various LeuT and MhsT structures in different conformation states, highlights that the interaction of this glutamate on TM2 with the unwound TM6 is maintained through the transport cycle (Figure S5). Given the importance of this interaction to transport, it is remarkable that in MhsT the significant movements of unwound TM6 are accommodated and play a crucial role in substrate specificity. The corresponding residue in other SLC6 transporters (Figure S2A) is to Glu84^DAT^, Glu136^SERT^ and Glu62^LeuT^. In most cases it interacts with another glutamate residue on TM10 (Glu490^DAT^, Glu508^SERT^, Glu419^LeuT^) and the interaction has been noted as important for protein stability and conformational changes [54]. The mutation of the interacting glutamate residue on TM10 to a lysine is one of the disease mutations causing Hartnup disease in SLC6A19 [55]. This invites the question why it is conserved as a glutamic acid and could not be replaced by e.g. a glutamine residue. Presumably, Glu66 (and its equivalent residue in other SLC6 transporters) is protonated and neutral in the buried environment (PROPKA estimates a pKa of 7.6 (molecule A) and 8.1 (molecule B) for the L-Val complex and 6.9 (partially protonated) for the Trp complex [56]), but the hydrogen-position of such a functionality is dislocated and can shift to support a rapid and dynamic change in hydrogen bonding capacities that promotes transport, unlike for a glutamine side chain that would have to rotate fixed hydrogen bonding geometries to align with dynamic transitions associated with transport.

Many transporters belonging to other SLC families share the LeuT fold despite of low sequence similarity, implying that structural as well as mechanistic similarities are present, e.g., the arginine/ agmatine antiporter, AdiC with arginine bound [19] and the L-5-benzyl-hydantoin bound sodiumbenzylhydantoin transporter, Mhp1 [20]. In both cases, the substrates are bound to the transporters in a similar manner substrates are bound to LeuT and MhsT, with an additional π-cation interaction in the case of AdiC and a π-stacking interaction in the case of Mhp1. Similarly, looking at the apo state of the organocation transporter, ApcT [21], a water filled cavity large enough to accommodate an L-Phe substrate molecule, is found at the same location. However, the betaine transporter, BetP [22], the galactose transporter, vSGLT [23]; and L-carnitine transporter, CaiT [24] transporters have binding pockets shifted towards TM2 and TM7, but with the TM1 and TM6 still participating in substrate binding. Notably, however, the conserved (G/A/C)ΦG sequence in TM6 of SLC6 transporters is not conserved within the unwound region of TM6 of transporters that do not belong to the SLC6 family, despite that fact that they share the overall LeuT fold. This is pointing to a specific role of an unwound region in TM6 in the case of SLC6 transporters including the NSS and AAT subfamilies. Furthermore, the conserved Glu residue in TM2 (Glu66 in MhsT) is only conserved among the SLC6 members, further corrobating the notion of distinct transport mechanisms along the lines of the existing gene families. The common LeuT fold therefore appears to represent an archaetype of transporters that, by rather small structural modifications, has evolutionarily developed into distinct transporter families with varying substrate translocation mechanisms for different physiological conditions and substrates.

## Materials and Methods

### Reagents

All chemicals were obtained from Sigma, unless stated otherwise. Amino acids stocks were prepared at 10 mM concentration in H_2_O and were stored at 4 °C. Nimesulide stock was prepared in DMSO and was stored at 4°C, protected from light.

### Protein expression and purification for structural studies

The *mhsT* gene was cloned into the pNZ8048 vector containing a chloramphenicol resistance gene, an N-terminal 1 Iis-tag and a TEV protease cleavage site. The protein was expressed in *Lactococcus lactis* NZ9000 strain and purified as described previously [33].

### Protein relipidation and crystallization

Purified MhsT, concentrated to approx. 9 mg/ml, was relipidated overnight in 1,2-dioleoyl-*sn*-glycero-3-phosphocholine (DOPC) at w/w ratio 3:04 and 3:08 protein:lipids, according to the method described previously [33]. Prior to crystallization experiment, protein sample was spun down at 290,000 x g for 20 min and mixed with either n-octyl-β-D-glucopyranoside (OG; Anatrace) or n-nonyl-β-D-glucoside (NG; Anatrace) detergent to final detergent concentration of 4 CMC.

Crystallization was set up at 19°C using vapor diffusion, hanging drop method, with immersion oil as a sealing agent. A reservoir buffer composition covered: 14-24 % PEG400, 0.3-0.5 NaCl, 100 mM Tris-HCl or HEPES-NaOH pH 7.0, 5 % or 10 % glycerol, 5 % Trimethylamine N-oxide (TMANO). Obtained crystals were harvested at 4°C and tested for diffraction at the Diamond synchrotron facility using beamline I24 and I04, Swiss Light Source using beamline PXI.

### Data processing and structure refinement

Obtained datasets were processed in P2 or P2_1_ space groups using XDS package [57] and CCP4 program suite [58]. The initial phases for the structures were obtained using Phaser [59] with MhsT-Trp structure (PDB ID: 4US3) [33] as a search model excluding TM5 from the model. Initial models were rebuilt manually in COOT [60] and atomic models were refined using phenix.refine [61]. The quality of the datasets and presence of twinning were checked in phenix.xtriage. The dataset for MhsT-Leu was refined using the (h -k -l) twin law. The final data and refinement statistics are presented in Table 1. Attached figures were prepared in PYMOL [62] the multiple sequence alignment was performed using MUSCLE [63] and the LOGO representation was obtained in CLC Main Workbench (CLC Bio, Qiagen).

### Protein expression for functional studies

Desired mutations were introduced into *mhsT* wild-type gene via site-directed mutagenesis (Mutagenesis QuikChange Lightning Kit, Agilent Technolgies, Inc) using designed primers. Final constructs were verified via DNA sequencing. Subsequently, the *mhsT* variants were cloned into pQO6TEV vector which is a modified version of pQE30, and were expressed in *E. coli* strain MQ614 [aroP mtr tnaB271::Tn5 tyrP1 pheP::cat] as described previously [33]. Briefly, an overnight preculture of *E. coli* cells was diluted in LB medium supplemented with 100 μg/mL ampicillin to an OD_420_ of 0.1. The culture was incubated at 37°C with shaking, until the OD reached 1 when the expression was induced by addition of 0.3 mM IPTG. Induced cells were incubated at 37 °C with shaking for 2 additional hours. Afterwards, the cells were harvested by centrifugation for 10 min at 13200 x g, at 4 °C. The pellet was washed twice in 100 mM Tris/MES, pH 7.5 and stored on ice until the uptake experiment.

### Transport measurements in intact E. coli cells

Uptake of L-[^3^H]-Trp (18 Ci/mmol) or L-[^3^H]-Leu (120 Ci/mmol; both American Radiolabeled Chemicals, Inc.) was measured in intact *E. coli* MQ614 [64] transformed with pQE60 or its derivatives harboring indicated MhsT variants. Cells were prepared for uptake studies as described [64] and uptake was performed in 10 mM Tris/Mes, pH 8.5, 150 mM NaCl at a final total cellular protein concentration of 0.035 mg/mL in the presence or absence of substrates or inhibitors as indicated. 100 μL samples were assayed for the indicated time periods and the uptake reactions were quenched by the addition of 100 mM KP_i_, pH 6.0, 100 mM LiCl. Cells were collected on Advantce MFS GF75 glass fiber filters. The accumulated radioactivity was determined (as counts per minute, cpm) in a Hidex 300 SL scintillation counter. Known amounts of radioactivity were used to determine the cpm-to-pmol conversion.

### Immunological protein detection

Relative amounts of the respective MhsT variants in the membrane of MQ614 were detected by Western blotting using a monoclonal antibody against the N-terminal His tag present in all MhsT constructs. 10 μg of total membrane protein in membrane vesicles of MQ614 harboring the indicated MhsT variant (or the control plasmid) were subjected to 11 % SDS-PAGE followed by incubation of the membrane with the His probe antibody (Santa Cruz Biotechnology, Inc.) and horseradish peroxidase-based chemiluminescence detection (SuperSignal^®^ West Pico kit, Thermo Scientific).

### Data Analysis

All uptake measurements were performed in duplicate or triplicate and repeated at least 5 times. Data (shown as mean ± S.E.M of triplicate determination) are from representative experiments in which all constructs were assayed in parallel. The Michaelis-Menten constant (*Km*) and maximum velocity (*V_max_)* of transport were determined by fitting the data of 10-second uptake measurements plotted as function of the concentration of the respective substrate to the Michaelis-Menten equation in GraphPad Prism 7.0. They are shown as mean ± S.E.M of the fit.

## Acknowledgements

The authors are grateful to technical assistance by Tetyana Klymchuk, Lotte T. Pedersen, Anna Marie Nielsen, and Audrey Warren and support for computing by Jesper L. Karlsen. We thank Steffen Sinning and Birgit Schiøtt for fruitful discussions. Work on the project was partially supported by a Short Term EMBO Fellowship [ASTF 80-2016] and Boehringer Ingelheim Fonds travel grant (2015) to D.F. The research was supported by NIH grants U54 GM087519 to JAJ, R01GM119396 to MQ, and by the Lundbeck Foundation grants 2012-12576 and 2016-2518 to PN.

## Author contributions

DF,CN: equal contributions on production of MhsT-substrate complexes and crystallization. DF, JL: data collection, processing and model refinement of MhsT-Tyr, MhsT-4FPhe, MhsT-Leu; CN: data collection, twinning analysis, processing and model refinement of MhsT-Tyr, MhsT-Phe, MhsT-Leu, MhsT-Val, and data processing, twinning analysis, and model refinement of MhsT-Ile with inputs from JL and PN; AEB: crystallization of MhsT-Ile, RB: expression, purification and initial crystallization of MhsT-Val; DF, IOS, MQ: E. coli uptake experiments with inputs from JAJ; LM: initial experiments on crystallization; PN: project design and supervision. Manuscript was drafted by DF and CN with input from JL, MQ, JL and PN.

## Supplementary

**Figure S1:** Examples of MhsT crystals obtained in complexes with different substrates. **1:** MhsT+Leu complex obtained using HiLiDe in hanging drop plates, 0.5 mM L-Leu, 0.1 M HEPES-NaOH pH 7.0, 16 % PEG 400. 0.4 M NaCl, 2x cmc OTG, 10 % glycerol; **2:** MhsT+4-F-Phe complex obtained using HiLiDe in hanging drop plates, 0.5 mM L-Phe, 0.1 M Tris-HCl pH 7.0, 14 % PEG 400, 0.4 M NaCl, 5 % TMANO, 2x cmc OTG, 5 % glycerol; **3:** MhsT+Val complex crystals obtained in hanging drop plates with protein:DOPC ration of 3:0.8 (w/w), 2x cmc NG; 5 % glycerol, 12 % PEG400, 0.1M HEPES-NaOH pH 7.0, 0.3 M NaCl, 5 % TMANO.

**Figure S2:** Conservation of the A) Glu66^MhsT^ and B) GMG motif within the unwound part of TM6 across the human SLC6 family membranes and their homologues from other organisms. Accession numbers: SLC6A1_Hs - P30531; SLC6A2_Hs - P23975; SLC6A3_Hs - Q01959; SLC6A4_Hs - P31645; SLC6A5_Hs - Q9Y345; SLC6A6_Hs - P31641; SLC6A7_Hs - Q99884; SLC6A8_Hs - P48029; SLC6A9_Hs - P48067; SLC6A11_Hs - P48066; SLC6A12_Hs - P48065; SLC6A13 - Q9NSD5; SLC6A14_Hs - Q9UN76; SLC6A15_Hs - Q9H2J7; SLC6A16_Hs - Q9GZN6; SLC6A17_Hs - Q9H1V8; SLC6A18_Hs - Q96N87; SLC6A19_Hs - Q695T7; SLC6A20_Hs - Q9NP91; SLC6A1_Rn - P23978; DAT_Dm - Q7K4Y6; SLC6A10_Sm - A0A0H3YEX6; TNAT_St - O50649; MJ1319_Mj - Q58715; METP_Cg - A0A0D6FTY1; MhsT - Q9KDT3; LeuT - O67854.

**Figure S3:** Binding of Trp (PDB ID: 4US3). Ligand binding site of MhsT in complex with L-Trp and volume of the binding pockets visualised by means of the van der Waals representation of the ligand and coordinating residues of MhsT in complex with L-Trp.

**Figure S4:** Binding site volumes calculated by use of voidoo for MhsT in complex with Phe, 4FPhe, Tyr, Val, Leu and Ile.

**Figure S5:** Movement of TM6, TM2 and Glu66 during the transport cycle. Structures are aligned based on scaffold TMs 3-4, and TMs 8-9.

LeuT outward open conformation (PDB ID: 3TT1): gray color; LeuT outward occluded conformation (PDB ID: 2A65) – orange color; Mhst-Val inward occluded conformation with a small substrate – green color; MhsT-4FPhe inward occluded conformation with an aromatic substrate-blue color; LeuT inward open conformation (PDB ID: 3TT3) – red color; LeuT return occluded conformation (PDB ID: 5JAE)-cyan color.

**Figure S6:** F_o_-F_c_ simulated annealing omit maps for MhsT in complex with L-Phe, L-4-F-Phe, L-Tyr and L-Val contoured at ± 3.0 r.m.s.d. Omit maps for MhsT in complex with L-Leu and L-Ile contoured at ± 2.5 r.m.s.d.

**Figure S7:** 2F_o_-F_c_ electron density maps contoured at 1.0 r.m.s.d. for the unwound region of TM6 of MhsT in complex with L-Phe, L-4-Phe, L-Tyr, L-Val, L-Leu and L-Ile.

**Figure S8:** A-F) F_o_-F_c_ simulated annealing omit maps of the water molecules near the GMG loop contoured at ± 3.0 r.m.s.d. of MhsT in complex with L-Phe, L-4-F-Phe, L-Tyr, L-Val, L-Leu and L-Ile.

**Figure S9:** Competitive uptake assay of selected amino acids in the presence of ^3^H-labeled L-leucine. MhsT-WT upake of L-leucine is inhibited to various degree by all of the hydrophobic amino acids. M236F mutant shows complete inability to compete L-luecine out with hydrophobic amino acids, indicating impaired binding or transport. Triple18 variant harbours following substitutions: L328T, M236F, V331A.

